# Proteomic profiling identifies Serpin B9 as mediator of resistance to CAR T-cell and bispecific antibody treatment in B-cell lymphoma

**DOI:** 10.1101/2023.06.26.546507

**Authors:** Berit J. Brinkmann, Tümay Capraz, Tobias Roider, Mareike Knoll, Carolin Kolb, Yi Liu, Antonia-Eugenia Angeli-Terzidou, Nagarajan Paramasivam, Björn Chapuy, Volker Eckstein, Tim Sauer, Michael Schmitt, Andreas Rosenwald, Carsten Müller-Tidow, Wolfgang Huber, Sascha Dietrich

**Affiliations:** Department of Medicine V, Hematology, Oncology and Rheumatology, University of Heidelberg, Heidelberg, Germany; Molecular Medicine Partnership Unit (MMPU), Heidelberg, Germany; European Molecular Biology Laboratory (EMBL), Heidelberg, Germany; Clinical Cooperation Unit Molecular Hematology/Oncology, German Cancer Research Center (DKFZ), Heidelberg, Germany; Faculty of Biosciences, University of Heidelberg, Heidelberg, Germany; Computational Oncology, Molecular Precision Oncology Program, National Center for Tumor Diseases (NCT), German Cancer Research Center (DKFZ), Heidelberg, Germany; Department of Hematology, Oncology and Cancer Immunology, Charité - University Medical Center, Berlin, Germany; Institute of Pathology, University of Würzburg, Würzburg, Germany; Department of Hematology and Oncology, University Hospital Düsseldorf, Düsseldorf, Germany

**Author notes:** **Corresponding author:** Sascha Dietrich, MD, Email address, Mailing address: Department of Hematology and Oncology, University Hospital Düsseldorf, Moorenstr. 5, 40225 Düsseldorf, Germany. These authors contributed equally to this work (Berit J. Brinkmann, Tümay Capraz, Tobias Roider). These senior authors contributed equally to this work (Wolfgang Huber, Sascha Dietrich).

## Abstract

Although T-cell-engaging therapies are highly effective in patients with relapsed and/or refractory B-cell non-Hodgkin lymphoma (B-NHL), responses are often not durable. To identify tumor-intrinsic drivers of resistance, we quantified *in-vitro* response to CD19-directed chimeric antigen receptor T-cells (CD19-CAR) and bispecific antibodies (BsAb) across 46 B-NHL cell lines and measured their proteomic profiles at baseline. Among the proteins associated with poor *in-vitro* response was Serpin B9, an endogenous granzyme B inhibitor. Knock-out of *SERPINB9* in cell lines with high intrinsic expression rendered them more susceptible to CD19-CAR and CD19-BsAb. Overexpression in cell lines with low intrinsic expression attenuated responses. Polatuzumab, vorinostat, lenalidomide, or checkpoint inhibitors improved response to CD19-CAR, although independently of Serpin B9 expression. Besides providing an important resource of therapy response and proteomic profiles, this study refines our understanding of resistance in T-cell engaging therapies, and suggests clinically relevant combination regimes.

## Introduction

Treatment options of B-cell non-Hodgkin lymphomas (B-NHL) now include bispecific antibodies (BsAb) and genetically engineered chimeric antigen receptor (CAR) T-cells targeting CD19 and CD20. Although such immunotherapies can induce durable responses in relapsed and refractory (r/r) diseases, long-term efficacy varies highly across patients. Depending on B-NHL entity and number of pre-treatments, 40-80 % of patients receiving CAR T-cells eventually progress or relapse within 12 months.^1–6^ An analogous situation with BsAb is currently being documented in clinical trials (e.g., NCT04712097, NCT04408638, NCT02811679). A better understanding of the mechanisms leading to unsustained or incomplete responses is needed for future improvements and potential stratification of T-cell-engaging therapies.

Both CAR T-cells and BsAb are able to elicit potent anti-tumor responses through induced T-cell activation. Accordingly, known response determinants include the composition and phenotype of transfused ^7–10^ as well as inherent T-cells ^11–13^. Besides, previous studies have identified few tumor cell-inherent resistance mechanisms, which predominantly involve loss or alterations of target antigens ^14–16^, or impairment of death receptor or interferon-γ signaling ^9,17–19^. These studies include large-scale loss of function screens in individual cell lines, which have established a fundamental understanding of resistance to CAR T-cell and BsAb therapy. Expanding to complementary approaches that better reflect and capture heterogeneity within and across B-NHL entities is urgently needed as a next logical step.

Here, we aimed to mimic heterogeneity within and across B-NHL entities by quantifying responses to CD19-CAR and BsAb of 46 established cell lines from diffuse large B-cell lymphoma (DLBCL), mantle cell lymphoma (MCL), Burkitt lymphoma (BL), follicular lymphoma (FL), and B-cell prolymphocytic leukemia (B-PLL). By correlating these response profiles with the cell lines’ protein expression profiles at baseline condition, we identified Serpin B9, an intracellular granzyme B (grB) inhibitor, as a key contributor to resistance to CAR T-cell and BsAb treatment. By CRISPR-mediated perturbation, we demonstrate that tumor-intrinsic levels of Serpin B9 strongly regulate sensitivity to T-cell-induced cytotoxicity. Combination of CD19-CAR with polatuzumab, vorinostat, lenalidomide, and checkpoint inhibitors improved response independently of Serpin B9 expression and without negatively impacting T-cell expansion.

## Results

### Cell line models of B-NHL respond heterogeneously to CAR T-cell and BsAb treatment

To capture the heterogeneity of B-NHL, we assembled a set of 46 established and authenticated cell lines derived from different B-cell lymphoma entities, including DLBCL (n = 24), MCL (n = 9), BL (n = 9), FL (n = 2), and B-PLL (n = 2). B-NHL cell lines were co-cultured for four days with different ratios of CAR T-cells (Figure 1A) or with non-transduced (NT) T-cells and bispecific antibodies targeting CD3 and CD19 or CD20 (CD19-or CD20-BsAb) at two different concentrations (Figure 1B). As control treatments, B-NHL cell lines were either cultured without any T-cells (mono-cultures) or co-cultured with NT T-cells. The percentage of viable lymphoma or T-cells after each treatment was measured based on the cell count normalized to the control treatment, NT T-cells without BsAb. NT T-cells did not induce considerable lymphoma cell killing, as was apparent when compared to mono-cultures (Supplementary Figure 1B). In contrast, both CD19-CAR and CD19-BsAb significantly killed lymphoma cells while T-cells expanded (Figure 1C).

**Figure 1.**
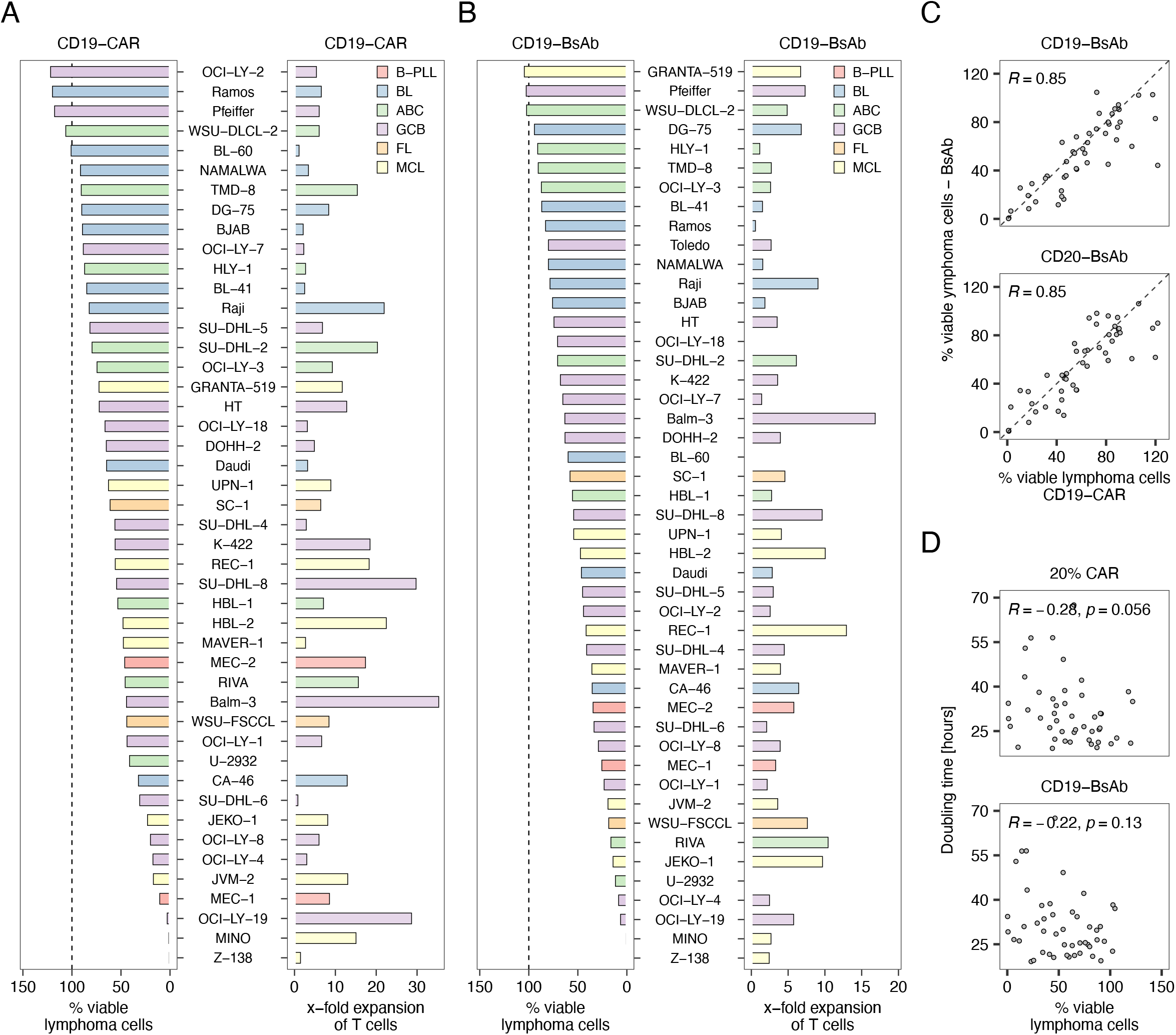
Response to treatment with CAR and BsAb is heterogenous across B-NHL cell lines. **A)** Percentage of viable lymphoma cells and **B)** x-fold expansion of T-cells after co-culturing B-NHL cell lines with CD19-CAR T-cells for four days. **C)** Percentage of viable lymphoma cells and **D)** x-fold expansion of NT T-cells after co-culturing B-NHL cell lines with NT T-cells in the presence of 10 ng/mL CD19-BsAb for for days. **E)** Spearman’s rank correlation of viable lymphoma cells after treatment with CD19-CAR and CD19-or CD20-BsAb. **F)** Scatter plot and spearman’s rank correlation of doubling time in hours and percentage of viable lymphoma cells after treatment with CD19-CAR and CD19-BsAb. NT: non-transduced. BsAb: bispecific antibody.

CAR- and BsAb-mediated killing of lymphoma cells was highly heterogeneous among the investigated cell lines but showed no association with the extent of T-cell expansion (Figure 1B, C). Killing of lymphoma cells correlated strongly between the different effectors, namely CD19-CAR, CD19-BsAb, and CD20-BsAb (Figure 1D). It increased with higher proportions of CAR T-cells or concentrations of BsAb (Supplementary Figure 1B). Comparing B-NHL entities, we found that BL responded significantly worse to CD19-CAR than MCL (Supplementary Figure 1D), which resembles clinically observed response patterns to CD19-CAR T-cells ^2,20,21^. Besides, neither proliferation activity, indicated as doubling time in hours during growth phase before treatment (Figure 1E), nor CD19 expression in lymphoma cells were associated with tumor cell killing (Supplementary Figure 1E), suggesting the involvement of B-cell inherent features in this heterogeneous response pattern.

### Proteomic profiling of B-NHL cell lines links Serpin B9 to impaired response to treatment with CAR and BsAb

To find B-cell-inherent factors driving the heterogeneous responses to CAR T-cells and bispecific antibodies, we used mass spectrometry to acquire whole cell proteomic profiles of all 46 cell lines, which revealed entity-specific differences (Supplementary Figure 2). Next, we looked for proteins whose abundance correlated with CD19-CAR or CD19-BsAb response (Figure 2A). We identified 30 proteins associated with CD19-CAR response and 107 with CD19-BsAb response (FDR = 0.05). We chose to follow up on Serpin B9 (Benjamini-Hochberg adjusted p-value 0.048 and 0.099 for CD19-CAR and CD19-BsAb, respectively), whose abundance was associated with poor response to both CD19-CAR and BsAb treatment. Serpin B9 is an endogenous inhibitor of granzyme B (grB) and has been described to regulate resistance to immune checkpoint inhibition in melanoma ^22,23^.

**Figure 2.**
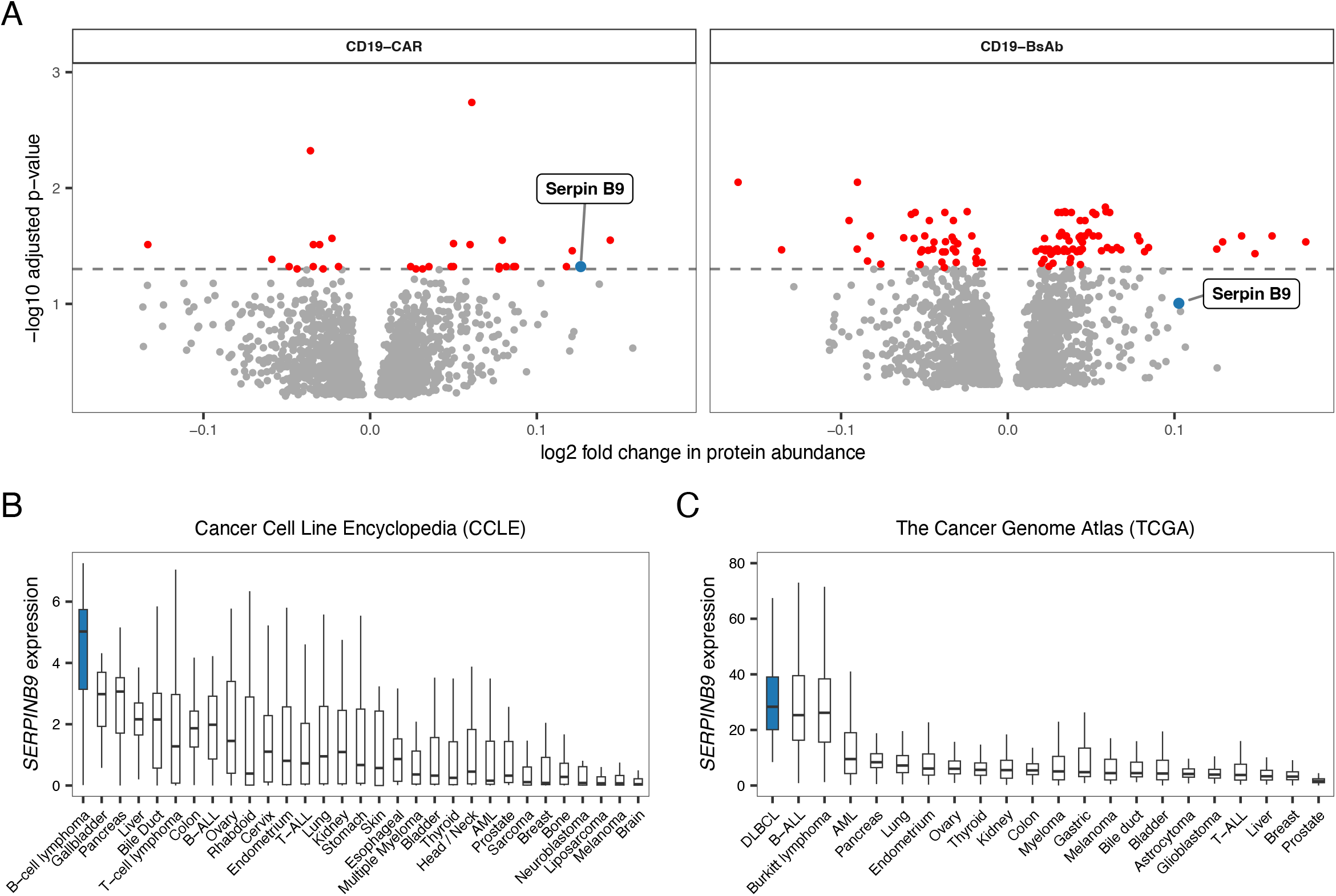
Proteomic profiling links Serpin B9 to impaired response to CAR and BsAb. **A)** Volcano plots showing log2 fold changes in protein abundances with respect to CD19-CAR and CD19-BsAb response. Proteins with high fold change are associated with a poor response to treatment. Adjusted p-values were calculated using the R package limma followed by multiple testing correction using the Benjamini Hochberg procedure (adjusted p-value = 0.05 indicated by the dashed line). **B)** *SERPINB9* expression across cell lines from various cancer entities (data from CCLE). **C)** *SERPINB9* expression in primary tumor samples across cancer types (data from TCGA). CCLE: Cancer Cell Line Encyclopedia. TCGA: The Cancer Genome Atlas.

This choice was corroborated by entity-specificity of its expression: Among the more than 1,000 cell lines of the Cancer Cell Line Encyclopedia (CCLE) ^24^, which represent a large range of cancer entities, we found that *SERPINB9* was most highly expressed in B-cell lymphoma (Figure 2B). Among the more than 10k primary tumor samples of The Cancer Genome Atlas (TCGA) ^25^ database, B-cell malignancies, including DLBCL, B-cell acute lymphoblastic leukemia (B-ALL) and Burkitt lymphoma, showed the highest levels of *SERPINB9* expression (Figure 2C).

### Knock-out of SERPINB9 increases susceptibility to CAR- and BsAb-mediated killing

To validate the proteomics data, we quantified Serpin B9 protein abundance by western blot in a representative set of 11 B-NHL cell lines (Supplementary Figure 3A), and observed good correlation (R = 0.87, Supplementary Figure 3B). In order to investigate if loss of Serpin B9 could reverse resistance to T-cell-mediated killing, we transduced DG-75 (BL), GRANTA-519 (MCL), and SU-DHL-5 (DLBCL) cells, which have comparably high intrinsic levels of Serpin B9 (Figure 3A), with a lentivirus delivering Cas9 and a guide RNA (gRNA) targeting *SERPINB9*. Loss of expression of *SERPINB9* (Sb9^KO^) was confirmed by western blot (Figure 3B). Similarly, we generated two stable and monoclonal cells lines overexpressing Serpin B9 by lentiviral transduction of CA-46 (BL) and OCI-LY-1 (DLBCL) (Sb9^OE^, Figure 3B), which have comparably low intrinsic protein levels (Figure 3A). Importantly, neither depletion nor overexpression of *SERPINB9* impacted on CD19 expression (Supplementary Figure 3C) or cell growth (Supplementary Figure 3D).

**Figure 3.**
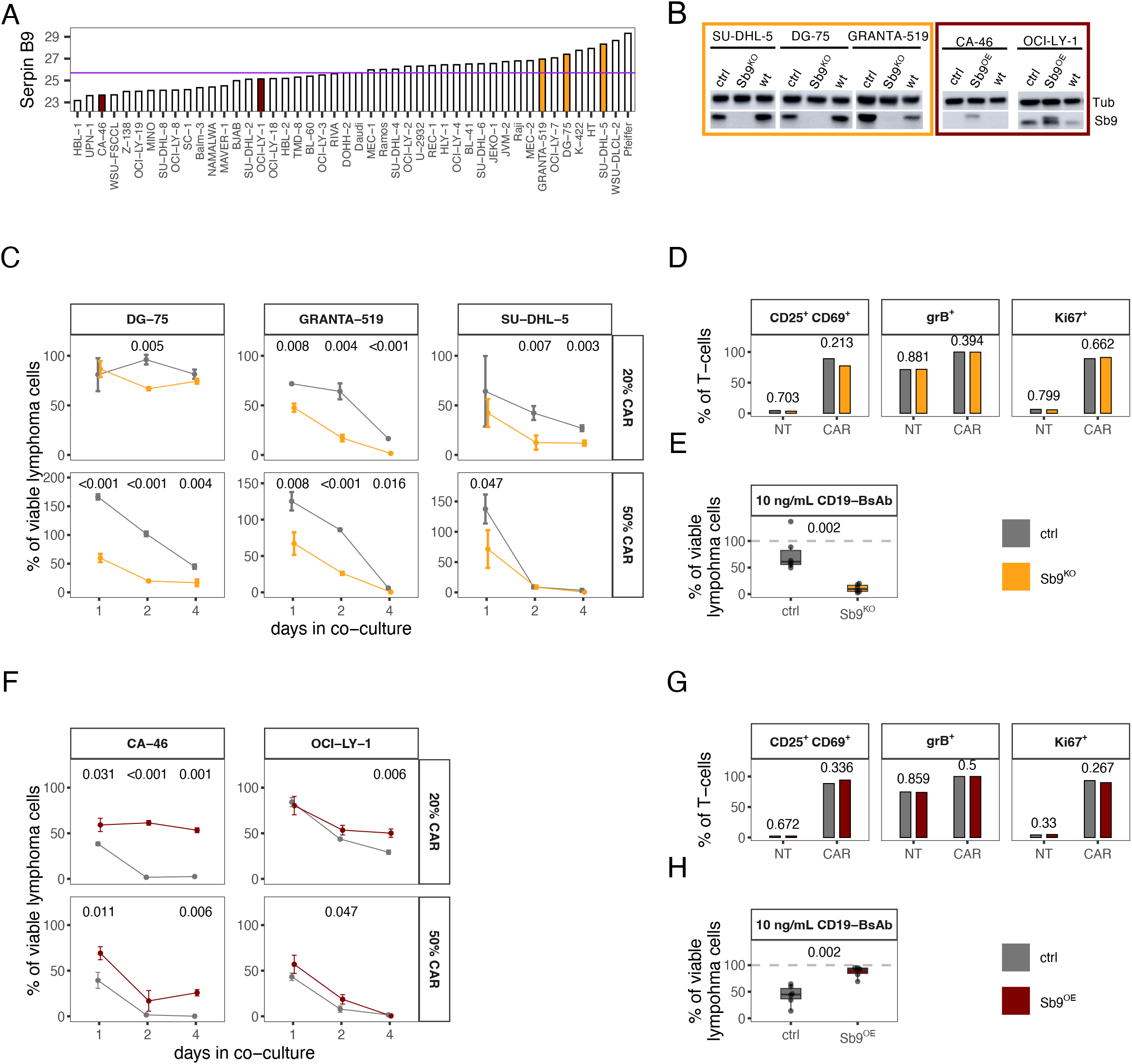
Serpin B9 regulates response to CAR and BsAb. **A)** Abundance of Serpin B9 in B-NHL cell lines. Yellow and red bars indicate cell lines with intrinsic Serpin B9 levels above or below median Serpin B9 expression (purple line) that were selected for knock-out or overexpression of *SERPINB9*, respectively. **B)** Western blot of Sb9^KO^, Sb9^OE^, ctrl, and wt cell lines. **C)** Percentage of viable Sb9^KO^ or ctrl cell lines after co-culture with indicated ratios of CD19-CAR T-cells for indicated incubation times. Values are shown as mean +/-SD of triplicates. **D)** Proportion of indicated T-cell subsets among NT or CAR T-cells co-cultured with Sb9^KO^ or ctrl cells. The mean of n=3 different B-NHL models is shown. **E)** Percentage of viable SU-DHL-5 Sb9^KO^ or ctrl cells after 3-day co-culture with PBMCs of n=6 different donors in the presence of 10 ng/mL CD19-BsAb. **F)** Percentage of viable Sb9^OE^ or ctrl cell lines after co-culture with indicated ratios of CD19-CAR T-cells for indicated incubation times. Values are shown as mean +/-SD of triplicates. **G)** Proportion of indicated T-cell subsets among NT or CAR T-cells co-cultured with Sb9^OE^ or ctrl cells. The mean of n=2 different B-NHL models is shown. **H)** Percentage of viable CA-46 Sb9^OE^ or ctrl cells after 3-day co-culture with PBMCs of n=6 different donors in the presence of 10 ng/mL CD19-BsAb. P-values comparing between Sb9^KO^ or Sb9^OE^ and ctrl cells were calculated using the two-sided t-test (C, D) or Wilcoxon’s test (E), and are denoted in the plots. Sb9: Serpin B9. Tub: α-Tubulin. KO: knock-out. OE: overexpression. ctrl: control. wt: wildtype. SD: standard deviation. grB: granzyme B. BsAb: bispecific antibody. PBMCs: peripheral blood mononuclear cells.

We measured response to CD19-CAR in Sb9^KO^ and control transduced cell lines and found that knock-out of *SERPINB9* rendered lymphoma cells more susceptible (Figure 3C). We further investigated if knock-out of *SERPINB9* affected the phenotype of co-cultured T-cells. Following three days of incubation, we detected high proportions of CD25^+^ CD69^+^, grB^+^, and Ki67^+^ cells among CAR T-cells, confirming an increased level of activation and proliferation compared to NT T-cells in our set-up (Figure 3D). Importantly, expression of activation and proliferation markers did not vary between T-cells co-cultured with Sb9^KO^ or with control cells (Figure 3D), suggesting that the observed differences in response were due to increased susceptibility of tumor cells rather than a change in T-cell phenotype.

To extend these results to BsAb treatment and thus to also rule out donor-specific effects, we co-cultured SU-DHL-5 Sb9^KO^ or control cells with peripheral blood mononuclear cells (PBMCs) from six different healthy donors in the presence or absence of CD19-BsAb. While on average only about 10 % of Sb9^KO^ cells remained viable, more than 60 % of the control cells survived treatment with BsAb (Figure 3E). Together, these results suggest that B-NHL cells with *SERPINB9* knock-out are more vulnerable to T-cell-mediated cytotoxicity, regardless of T-cell donor and treatment approach.

### Overexpression of SERPINB9 renders B-NHL cells more resistant to CAR and BsAb treatment

Next, we compared CAR T-cell-mediated killing of Sb9^OE^ and control transduced cell lines. Overexpression of *SERPINB9* rendered cells more resistant to CAR T-cell-mediated killing across different incubation periods and CAR T-cell ratios (Figure 3F). As for the Sb9^KO^ models, we characterized the phenotype of CAR and NT T-cells after co-culture with Sb9^OE^ and control cells. The proportion of CD25^+^CD69^+^, grB^+^, and Ki67^+^ cells among CAR T-cells co-cultured with Sb9^OE^ cells was not increased compared to those co-cultured with control transduced target cells (Figure 3G), demonstrating that the observed increase in killing of Sb9^OE^ cells was not mediated by altered T-cell activation or proliferation.

To investigate whether the observed effects were comparable for treatment with BsAb, we co-cultured CA-46 Sb9^OE^ and control cells with PBMCs of six healthy donors in the presence or absence of CD19-BsAb. On average, more than 50 % of the control transduced cells were killed by treatment with CD19-BsAb, whereas more than 85 % of Sb9^OE^ cells survived (Figure 3H). We conclude that Serpin B9 renders B-NHL cells more resistant to T-cell-mediated cytotoxicity, irrespective of T-cell donor and treatment approach.

### Pharmacological strategies to improve response to T-cell-mediated cytotoxicity

After identifying Serpin B9 as mediator of resistance to T-cell-mediated cytotoxicity, we aimed to determine combinatorial drug treatments that could improve poor response. To this end, we combined treatment with CD19-CAR with a panel of 11 clinically relevant drugs, including immune checkpoint inhibitors, antibody-drug conjugate polatuzumab vedotin, chemotherapeutics, and small molecules. We presumed that combination compounds of interest should increase killing of lymphoma cells while not deteriorating activation and proliferation of CD19-CAR. For this purpose, we quantified the change in CAR T-cell expansion (Figure 4A) and lymphoma cell killing (Figure 4B) mediated by the combination compound and normalized it to the corresponding condition with CD19-CAR only. A paired, two-sided Wilcoxon’s test was used to determine significant differences between treatment with CD19-CAR alone and in combination with a drug. We ranked the tested drugs by their impact on CAR T-cell expansion and found that the immunomodulatory drug lenalidomide and checkpoint inhibitors nivolumab and ipilimumab significantly improved not only expansion of CD19-CAR but also killing of lymphoma cells. Antibody-drug conjugate polatuzumab vedotin and histone deacetylase (HDAC) inhibitor vorinostat substantially increased killing of lymphoma cells while not affecting CD19-CAR expansion. Some of the remaining drugs, such as bendamustine, were highly toxic to lymphoma cells but also impaired T-cell expansion.

**Figure 4.**
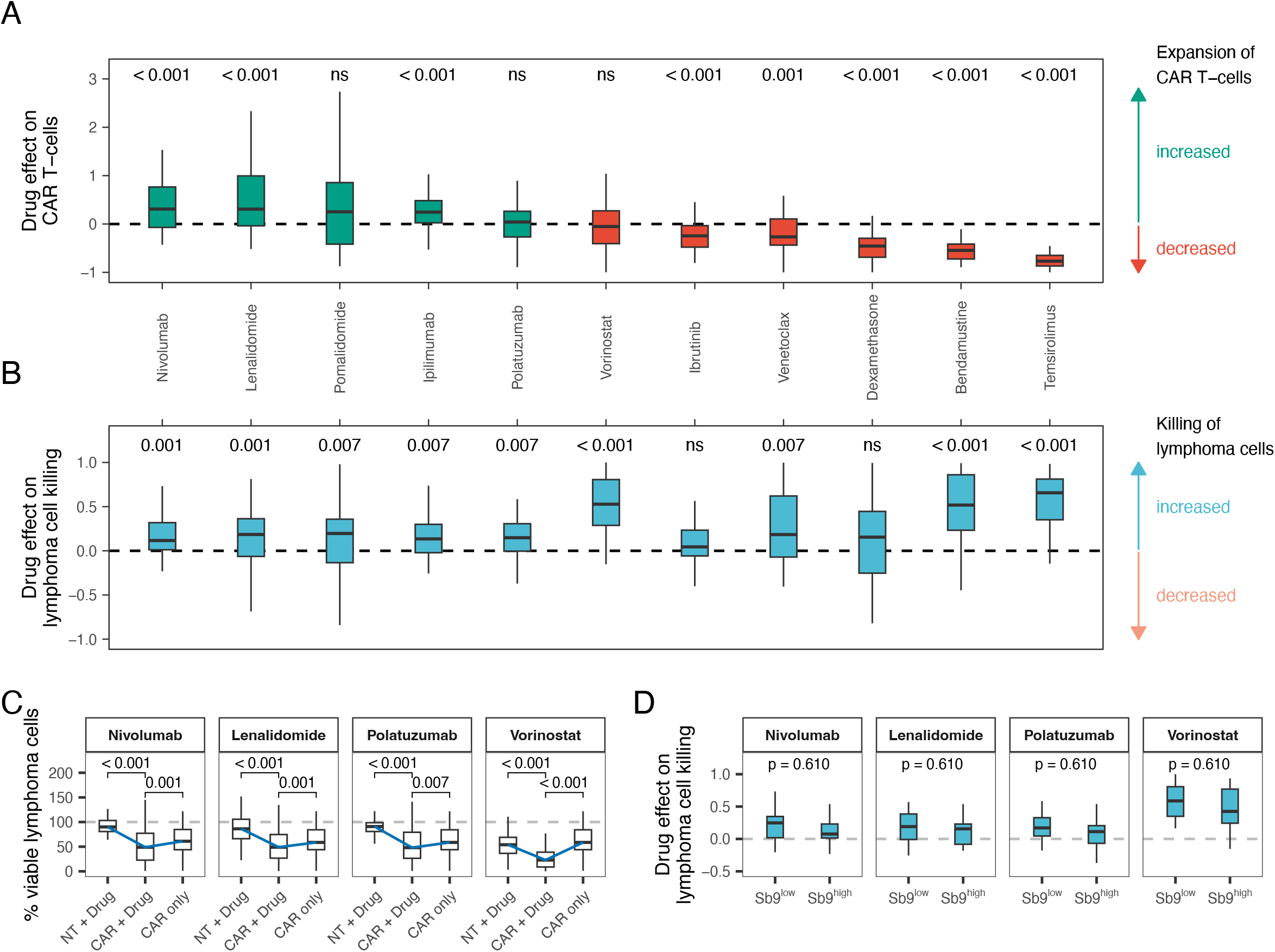
Drug-induced enhancement of CAR T-cell cytotoxicity is independent of Serpin B9. 46 B-NHL cell lines were co-cultured for four days with 20 % NT or CD19-CAR T-cells in the presence or absence of indicated drugs. **A, B)** Drug effects on CAR T-cell expansion and lymphoma cell killing were calculated as described in the methods section. Drugs are arranged in descending order by their effect on CAR T-cell expansion. **A)** Drug effects on CAR T-cell expansion. Green boxes mark drugs with a median increasing effect on CAR T-cell expansion, red boxes mark drugs with a median decreasing effect on CAR T-cell expansion. **B)** Drug effects on lymphoma cell killing. Blue boxes mark drugs with a median increasing effect on lymphoma cell killing. P-values from the comparison of CAR T-cell expansion (A) or lymphoma cell killing (B) with or without a given drug were calculated using the two-sided paired Wilcoxon’s test and are denoted in the plot. **C)** Percentage of viable lymphoma cells after 4-day co-culture with NT or CD19-CAR T-cells with or without a given drug. Blue lines connect the medians of each co-culture/drug-condition to ease interpretation. **D)** Comparison of drug effects on lymphoma cell killing between cell lines with Serpin B9 levels above (Sb9^high^) or below (Sb9^low^) median Serpin B9 abundance. P-values of indicated comparisons were calculated using the two-sided paired (C) or non-paired (D) Wilcoxon’s test and are denoted in the plots. ns: p-value > 0.05. Sb9: Serpin B9. NT: non-transduced.

We next assessed whether the combination of CD19-CAR and a given drug was more effective at killing lymphoma cells than the single treatments (Figure 4C, Supplementary Figure 4A). The combination of CD19-CAR and vorinostat, lenalidomide, polatuzumab, or immune checkpoint inhibitors killed lymphoma cells more effectively than any of the individual treatments (Figure 4C, Supplementary Figure 4A). None of these drugs negatively impacted T-cell expansion (Figure 4C, Supplementary Figure 4A). Comparing the increase in lymphoma cell killing by any given drug between cell lines with high or low Serpin B9 expression revealed no significant difference (Figure 4D, Supplementary Figure 4B), indicating that lymphoma cell killing was improved in general and independent of Serpin B9 protein abundance. Together, these results highlight that T-cell-mediated killing of lymphoma cells could be improved and given additional toxicity by drugs that either enhance or not disturb T-cell expansion. This drug-mediated improvement was not specific to samples with high Serpin B9 expression.

## Discussion

In this study, we quantified *in-vitro* responses to CD19-directed CAR T-cells, and CD19-or CD20-directed BsAb across a comprehensive set of 46 B-NHL cell lines. Response to CD19-CAR and BsAb was highly correlated, emphasizing their shared mechanism of action, T-cell-mediated cytotoxicity. By correlating response with proteomic profiles, we found that the abundance of Serpin B9, an endogenous grB inhibitor ^26^, was associated with reduced response to treatment with CAR T-cells and BsAb. We established a causality relationship using overexpression and knock-out models, which showed that Serpin B9 mediates resistance to CAR T-cell and BsAb treatment. Physiologically, Serpin B9 is highly expressed by cytotoxic lymphocytes and natural killer cells to protect them from misdirected grB ^26,27^. In tumor cells, Serpin B9 expression has previously been linked to poor outcome following immune checkpoint blockade in melanoma ^22,28^. In models of solid cancers, Serpin B9 was identified to protect tumor cells from T-cell-mediated cytotoxicity following irradiation via induction of type I interferon signaling ^29^. A recent study discovered a link between Serpin B9 expression and reduced killing of tumor cells by CAR T-cells in the DLBCL cell line OCI-LY-7 ^30^. Here, we identified Serpin B9 in an unbiased proteomic approach across a large number of heterogeneous B-NHL models as mediator of resistance not only to CAR T-cells but also to BsAb. Importantly, we demonstrate that protection mediated by Serpin B9 is present in different B-NHL entities and across different T-cell donors. Our results strengthen and extend these hints from the literature and identify Serpin B9 expression as an important tumor cell-intrinsic mechanism of resistance to T-cell engaging therapy. Future studies will need to confirm the prognostic role of Serpin B9 in B-NHL patients treated with CD19-CAR or BsAb.

In order to improve response to CAR T-cells, particularly in cell lines with high Serpin B9 expression, we combined clinically relevant drugs with CD19-CAR. The flow cytometry-based read-out enabled us to identify drugs that exert combination effects in different ways: Lenalidomide, nivolumab, and ipilimumab enhanced killing of lymphoma cells by promoting T-cell expansion, whereas polatuzumab and vorinostat exerted additional cytotoxicity. All of the identified drugs were similarly effective, regardless of the level of Serpin B9 expression in the target cells. We and others have previously demonstrated that lenalidomide and checkpoint inhibitors act synergistically on autologous lymph node-derived T-cells when combined with BsAb ^12,31^, underlining their potential as combination partners for T-cell engaging therapies. The combination of HDAC inhibitor vorinostat and CD19-CAR conveyed superior tumor cell-directed toxicity than either of the individual treatments alone. This additive effect could be explained by the fact that HDAC inhibitors are able to induce apoptosis ^32^ and have previously been demonstrated to render tumor cells susceptible to death receptor-induced apoptosis ^33^. Indeed, Torres-Collado *et al*. found that pre-treatment with vorinostat was able to re-sensitize B-NHL cell lines that had been resistant to death receptor-mediated killing by CAR T-cells ^34^. Besides vorinostat, polatuzumab vedotin provided lymphoma-specific toxicity owing to its ability to specifically recognize CD79b.

Taken together, the discovery of Serpin B9 as a mediator of resistance to T-cell engaging therapy extends our understanding of tumor-intrinsic resistance mechanisms to immunotherapy in B-NHL. By performing whole-cell proteomics in a large collection of B-NHL cell lines, we were able to capture heterogeneity across entities and additionally provide proteomic and response profiles as a valuable resource for the investigation of lymphoma biology. Together with the proposed combination drugs, our results may contribute to the optimization of CAR T-cell and BsAb treatments in B-NHL.

## Acknowledgements

T.R. was supported by a fellowship of the German Federal Ministry of Education and Research (BMBF) and a physician scientist fellowship of the Medical Faculty of University Heidelberg. S.D. was supported by a grant of the Hairy Cell Leukemia Foundation, the Heidelberg Research Centre for Molecular Medicine (HRCMM) and an e:med junior group grant of the German Federal Ministry of Education and Research (BMBF). S.D and B.C. were supported by the Gilead Research Scholar Program. We thank the FACS core facility of the University Hospital Heidelberg with Dr. Volker Eckstein, the Flow Core Facility of the EMBL and the Proteomics Core Facility (PCF) of the EMBL Heidelberg with Per Haberkant and Frank Stein for their excellent (technical) assistance. Results presented in Figure 2C and D are based upon data generated by the TCGA Research Network: https://www.cancer.gov/tcga.

## Author contributions

Conceptualization, S.D.; Data curation, T.R, T.C., Formal Analysis, B.J.B., T.R., T.C., N.P.; Funding acquisition, S.D.; Investigation, B.J.B, T.R., M.K., C.K., Y.L.; Methodology, B.J.B, T.R., A.-E.A.-T.; Resources, S.D., W.H., C.M.T., B.C., M.S., A.R., T.S., V.E.; Supervision, S.D., W.H.; Visualization, B.J.B., T.R., T.C.; Writing – original draft, B.J.B, T.R., T.C., W.H., S.D.

## Material and Methods

### Cell lines

Cell lines were cultured in RPMI, IMDM, DMEM or alpha-MEM (all Thermo Fisher Scientific) supplemented with 10-20% fetal bovine serum (FBS; Thermo Fisher Scientific), 1 % penicillin/streptomycin (Thermo Fisher Scientific) and 1 % glutamine (Thermo Fisher Scientific) in a humidified atmosphere at 37 °C and 5 % CO_2_. Cell lines were either purchased from DSMZ or kindly provided by Björn Chapuy. A detailed overview of the cell lines, medium composition and source is given in Supplementary Table 1. All cell lines were tested negative for mycoplasma.

### CAR T-cell manufacturing

Gamma-retroviral vector production and transduction of T-cells was performed as previously described^35^. Briefly, HEK 293T cells were co-transfected with the CD19-CAR encoding plasmid (kind gift of Prof. Malcolm Brenner from Baylor College of Medicine in Houston, Texas), Peg-Pam-e and RDF plasmid (both kindly provided by Prof. Michael Schmitt from the University Hospital Heidelberg), using GeneJuice (#70967-3, Sigma-Aldrich). Viral supernatant was collected two days post transfection. PBMCs were prepared by Ficoll density centrifugation of peripheral blood of healthy donors. Wells of a 24-well plate were coated with 1 μg/mL anti-CD3 (#317326) and anti-CD28 antibodies (#302934, both BioLegend). For activation of T-cells, PBMCs were seeded in AIM-V medium (#1205508, Thermo Fisher Scientific) supplemented with 10 % FBS, gentamicin/streptomycin and 1 % glutamine (Thermo Fisher Scientific). Activated T-cells were kept in medium supplemented with IL-7 and IL-15 (#200-07 and #200-15, PeproTech, 10 and 5 ng/mL, respectively). Gamma-retroviral transduction was facilitated by RetroNectin (#T100B, Takara Bio). Transduced and non-transduced (NT) control T-cells were expanded up to a maximum of 14 days. Transduction efficiency was measured by flow cytometry, staining with biotinylated Protein L (#29997, Thermo Fisher Scientific) and Streptavidin-PE (#405203, BioLegend).

### Flow cytometry-based screening

384-well plates (#781904, Greiner Bio One) were printed with 100 nL of solvent control or CD19-BsAb (1 ng/mL and 10 ng/mL, blinatumomab provided by the pharmacy of the University Hospital Heidelberg) and CD20-BsAb (100 ng/mL and 1,000 ng/mL, Genentech via material transfer agreement) using an automated pipetting system (CyBio Well vario, Jena Analytik). Two days before performing the screen, cell lines were cultured at 2.5 × 10^6^/mL in AIM-V medium (Thermo Fisher Scientific) supplemented with 10 % FBS, gentamicin/streptomycin and 1 % glutamine (Thermo Fisher Scientific). Doubling time during this phase was calculated by comparing cell counts from day 0 and 2. For labeling, target cells were incubated with 2 µM CellTracker Green (#C2925, Thermo Fisher Scientific) for 30 minutes, according to manufacturer’s instructions. Per well, either 4,000 or 8,000 cells were seeded using the CyBio Well vario, depending on whether the cells were slow (doubling time > 30 h) or fast growing (doubling time < 30 h). CAR or NT T-cells were added to solvent control wells to make up 10 and 20 % of the total number of cells per well, NT T-cells were added to BsAb-containing conditions to make up 20 % of the total number of cells per well. Following four days of incubation, cells were stained with fixable viability dye eFluor 506 (#65-0866-18), anti-CD3-PE (#300308) and anti-CD19-APC (#302212, all BioLegend), using the CyBio Well vario. Finally, CountBright Absolute Counting Beads (#C36950, Thermo Fisher Scientific) were added before samples were acquired on an LSR Fortessa equipped with an HTS system and BD FACSDiva software (BD Biosciences).

### Analysis of flow cytometry data

Obtained flow cytometry data were gated using FlowJo (BD Biosciences). An exemplary gating strategy to identify viable lymphoma, CAR, and NT T-cells is shown in Supplementary Figure 5A. Absolute numbers of lymphoma and T-cells were calculated based on the acquired count of counting beads, according to manufacturer’s instructions. Obtained lymphoma cell counts were divided by the mean absolute number of lymphoma cells in control co-cultures with NT T-cells and multiplied by 100 % in order to calculate the mean percentage of viable lymphoma cells. X-fold expansion of T-cells was calculated by dividing the obtained cell counts of T-cells by the mean absolute number of T-cells in control co-cultures with NT T-cells.

### Proteomics profiling and data processing

A detailed description of methodology used for proteomic profiling is provided in the Supplementary Methods. Briefly, pellets of 5×10^6^ cells per B-NHL cell line were washed two times with PBS and snap-frozen. Cell pellets lysed by 1 % SDS lysis buffer and further prepared using a modified version of the Single-Pot Solid-Phase-enhanced Sample Preparation (SP3) protocol ^36,37^. Peptides were labeled with TMT10plex (Thermo Scientific) label reagent and purified by a reverse phase clean-up step (OASIS HLB 96-well µElution Plate, Waters)^38^. Purified peptides were subjected to an offline fractionation under high pH conditions ^37^. Resulting fractions were then analyzed by LC-MS/MS on an Orbitrap Fusion Lumos mass spectrometer (Thermo Scientific) as previously described ^39^. Acquired data were analyzed using IsobarQuant ^40^ and Mascot V2.4 (Matrix Science) using a reverse UniProt FASTA Homo sapiens database (UP000005640 from May 2016). The raw output files of IsobarQuant (protein.txt-files) were processed using the R programming language (version 4.2.2). Preprocessing steps were done using the R package matrixQCvis ^41^ (version 1.4.0). Only proteins that were quantified with at least two unique peptides and present in at least 6 TMT channels were considered for the analysis. Raw signal-sums (signal_sum columns) were normalized using variance stabilization normalization^42^ followed by MinDet imputation, setting each missing protein abundance to their 0.01 quantile.

### Protein differential abundance analysis

Protein differential abundance between cell lines (n_samples_ = 70 and proteins = 4873) with respect to their percentage of viable lymphoma cells after treatment with CAR T-cells and BsAb, was assessed using the R package limma ^43^ (version 3.52.4). To account for potential batch effects TMT channels were included as covariates. P-values were adjusted using the Benjamini-Hochberg procedure ^44^. Differentially abundant proteins were defined as adjusted p-value < 0.05.

### Establishment of SERPINB9 knock-out cells

*SERPINB9* was deleted via CRISPR/Cas9. Guide ribonucleic acids (gRNAs) were designed using Off-Spotter^45^. In total, four different gRNAs (two targeting exon 1, one targeting exon 2 and one targeting exon 4) with the highest score of inverse likelihood of off-target binding were selected. One scramble control gRNA was designed with no targeting site within the genome. gDNAs were synthesized by Biolegio (Biolegio, Nijmegen, Netherlands). Sense and antisense-strand gDNAs were denatured at 95 °C for 10 min, followed by stepwise annealing. Annealed double stranded DNA were inserted into vector pL-CRISPR.EFS.GFP (addgene plasmid #57818) digested with restriction enzyme Esp3I (Thermo Fisher Scientific). Correct clones were confirmed by Sanger sequencing. Lentiviruses were produced as described previously. ^46^ In brief, plasmid pL-CRISPR.EFS.GFP-gDNA was co-transfected with containing lentiviral packaging plasmids pLP1, pLP2 and pLP/VSV-G into HEK 293T cells using TurboFect (#R0531, Thermo Fisher Scientific). Lenti-virions were collected after 3 days and concentrated by ultracentrifugation at 4 °C. Transduction of B-NHL cell lines was performed via spinoculation using Polybrene (#TR-1003-G, Sigma-Aldrich). Successfully transduced cells were selected by FACS. Based on GFP expression, single cells of SU-DHL-5 and DG-75 were sorted to obtain monoclonal populations. Depletion of Serpin B9 was verified by western blotting. gRNA #2 (5’-CCTGGCCATGGTTCTCCTAG-3’) successfully depleted Serpin B9.

### Establishment of SERPINB9 overexpression cell lines

To generate lentiviral particles, a lentiviral expression vector containing a turbo green fluorescent protein (turboGFP) reporter and the open reading frame of *SERPINB9* or, as control, turbo red fluorescent protein (turboRFP) (Precision LentiORF, Horizon Discovery) were co-transfected with psPAX2 (addgene plasmid #12260) and pMD2.G (addgene plasmid #12259) into HEK 293T cells using TurboFect (Thermo Fisher Scientific). Viral supernatant was collected on day 3 and 4 after transfection, filtered through a 0.45 µm filter, pooled and concentrated by ultracentrifugation at 4 °C. Viral particles were resuspended in ice cold PBS and stored at -80 °C until use. Transduction was performed by spinoculation using Polybrene (#TR-1003, Sigma-Aldrich). Successfully transduced cells were selected using blasticidin. To obtain a monoclonal population, single cells were sorted based on GFP expression. Overexpression of Serpin B9 was confirmed by western blotting.

### Western blotting

Cell pellets were lysed in RIPA buffer (Sigma-Aldrich). 20 µg of lysates were run on 10 % acrylamide gels and transferred to PVDF membranes. Primary antibodies of Serpin B9 (Santa Cruz, #sc-57531) and α-Tubulin (#66009-1-Ig, Proteintech Group,) and the secondary antibody anti-mouse-IgG-HRP-conjugated (#SA00001-1,Proteintech Group) were used for detection. To quantify protein expression, band intensities were quantified and normalized to the internal control.

### Flow cytometry of Sb9^KO^ and Sb9^OE^ cell lines

Sb9^KO^ and Sb9^OE^ cell lines were labeled as described above and co-cultures of 35,000 cells were seeded per well in 96-well plates, containing 20 % or 50 % CD19-CAR or NT T-cells. For experiments with BsAb, 17,500 SU-DHL-5 Sb9^KO^ and CA-46 Sb9^OE^ cells were co-cultured with 35,000 PBMCs of six different donors with or without 10 ng/mL CD19-BsAb. Following one, two, and four days of incubation (in case of CD19-CAR co-cultures) and three days of incubation (in case of BsAb), staining was done using anti-CD19-BV421 (#302234), anti-CD3-APC (#300312), and viability dye e506 (#65-0866-18, all BioLegend). For quantification of activation markers, co-cultures with 50 % T-cells were collected on day three and stained with fixable viability dye e506 (#65-0866-18), anti-CD3-PerCP-Cy5.5 (#317336), anti-CD4-APC (#300514), anti-CD8-APC-Cy7 (#300926), anti-CD69-BV711 (#310944, all BioLegend). Intracellular staining was done using the fixation/permeabilization buffer (#00-5123-43 and #00-8333, Thermo Fisher Scientific) and anti-Ki67-BV786 (#563756m BD Biosciences) and anti-granzyme B-PE-Dazzle (#372216, BioLegend). Cells were acquired on a FACSymphony (BD Biosciences). An exemplary gating strategy to identify CD25^+^ CD69^+^, grB^+^, and Ki67^+^ cells is shown in Supplementary Figure 5B.

### Growth curves of Sb9^KO^ and Sb9^OE^ cell lines

A total of 500,000 Sb9^KO^, SB9^OE^, or control transduced cells were seeded in triplicates per well of a 6-well plate. Cells were cultured for four consecutive days and counted every day. Fresh medium was added if necessary.

### Drug-mediated modulation of CAR T-cell cytotoxicity

Cell lines were co-cultured with 20 % CD19-CAR as for flow cytometry-based screening but in the presence of 100 nL of a given drug or solvent control. Drugs and final concentrations used are provided in Supplementary Table 2. In order to quantify the change in T-cell expansion and lymphoma cell killing as illustrated in Figure 4A and B, drug effects were calculated as follows:

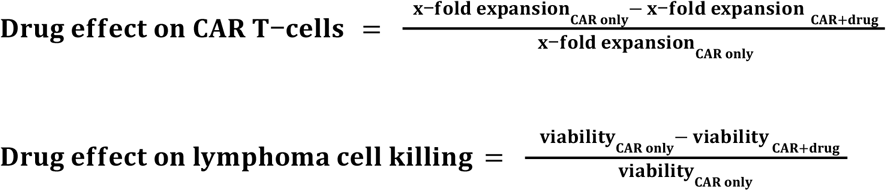

Using the two-sided paired Wilcoxon’s test, we tested for significant differences in CAR T-cell expansion (Figure 4A) and lymphoma cell killing (Figure 4B) with or without additional drug perturbation.

## Data availability

Proteomic data will be available in the PRIDE database (https://www.ebi.ac.uk/pride/) upon publication.

## Code availability

The computational codes, in the form of Rmarkdown documents, for reproducing all main and supplementary figures will be available upon publication.

