## Supplementary Figures for "Proteomic profiling identifies Serpin B9 as mediator of resistance to CAR T-cell and bispecific antibody treatment in B-cell lymphoma"

### Supplementary Figure 1

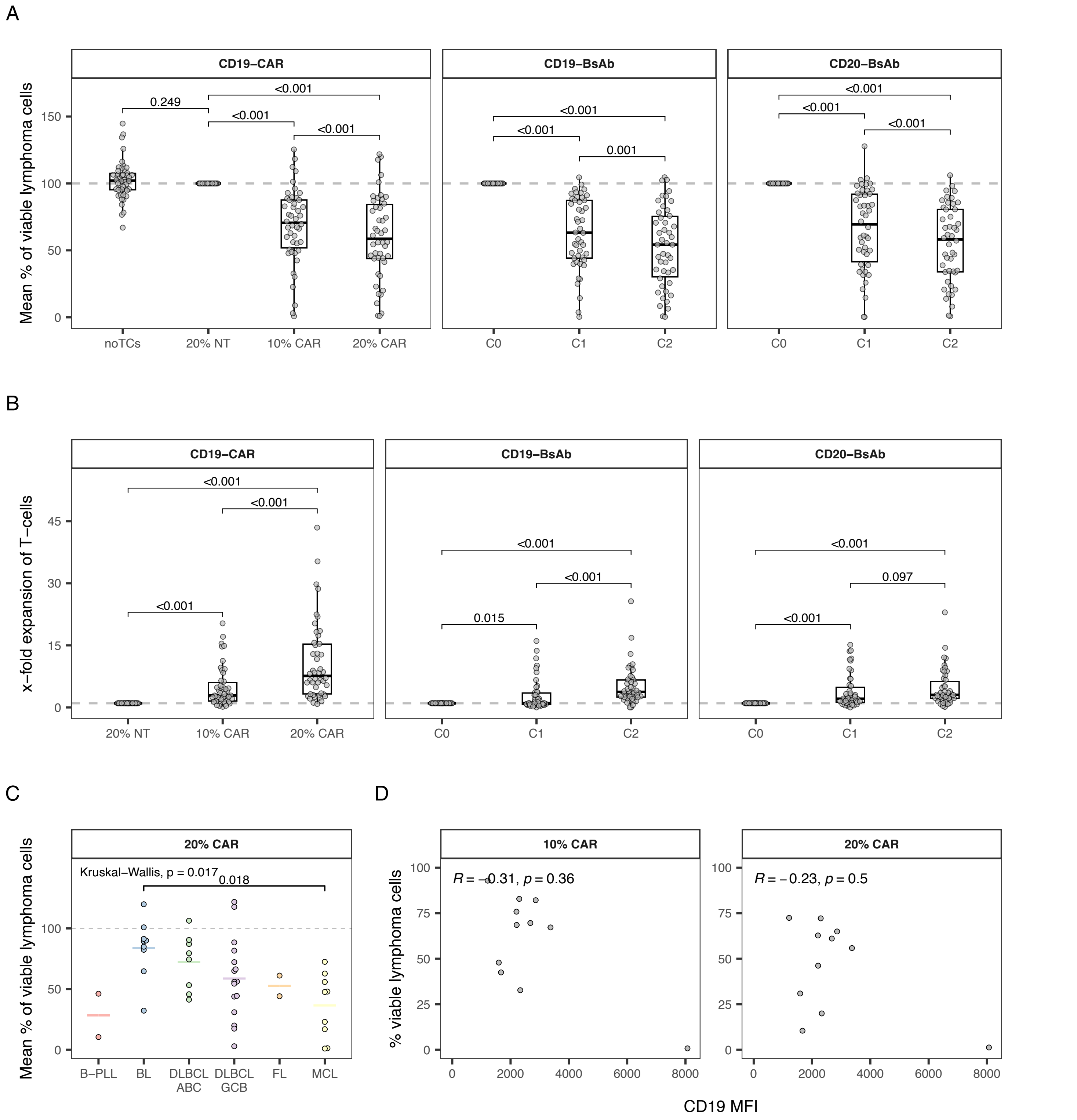


**Supplementary Figure 1. A)** Mean percentage of viable lymphoma cells after treatment with different ratios of CD19-CAR, NT T-cells or no added T-cells (left panel), treatment with with NT T-cells and different concentrations of CD19-BsAb (middle panel) or CD20-BsAb (right panel). **B)** X-fold expansion of NT and CD19-CAR T-cells after co-culture with B-NHL cell lines (left panel) and of NT T-cells after co-culture with B-NHL cell lines in the presence or absence of CD19-BsAb (middle panel) and CD20-BsAb (right panel). Dots indicate means per cell line. P-values were calculated between co-cultures with NT T-cells and every other condition using the paired, two-sided Wilcoxon's test. **C)** Comparison of the mean percentage of viable lymphoma cells after co-culture with 20 % CD19-CAR T-cells across different B-NHL entities. Entities were compared using the Kruskal-Wallis test and p-values were calculated post-hoc using the two-sided Wilcoxon's test. Points represent single cell lines. Horizontal bars represent mean values per entity. **D)** Spearman correlation's correlation of CD19 MFI in selected B-NHL cell lines and the percentage of viable cells after treatment with indicated ratios of CD19-CAR T-cells. NT: non-transduced. BsAb: bispecific antibody. C0: no BsAb. C1: low BsAb concentration. C2: high BsAb concentration.

### Supplementary Figure 2

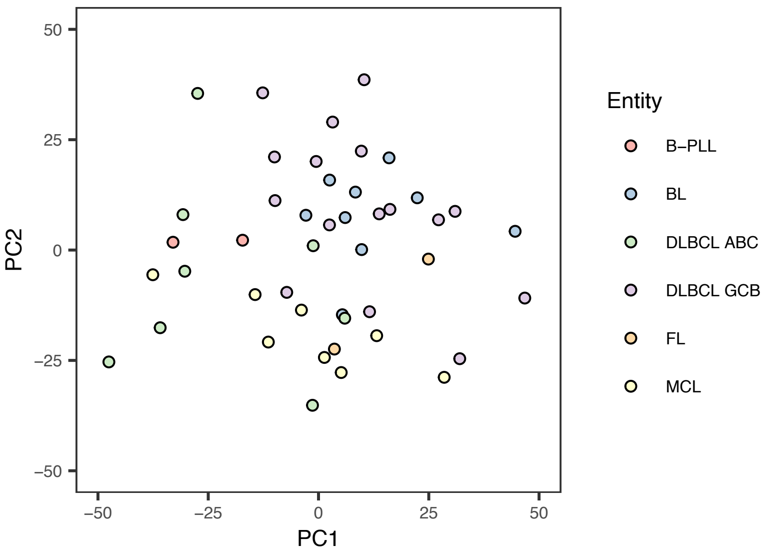


**Supplementary Figure 2.** Scatterplot of first the two principal components of the proteomics data of all 46 B-NHL cell lines (proteins = 4873). Each B-NHL cell line is colored according to their entity.

### Supplementary Figure 3

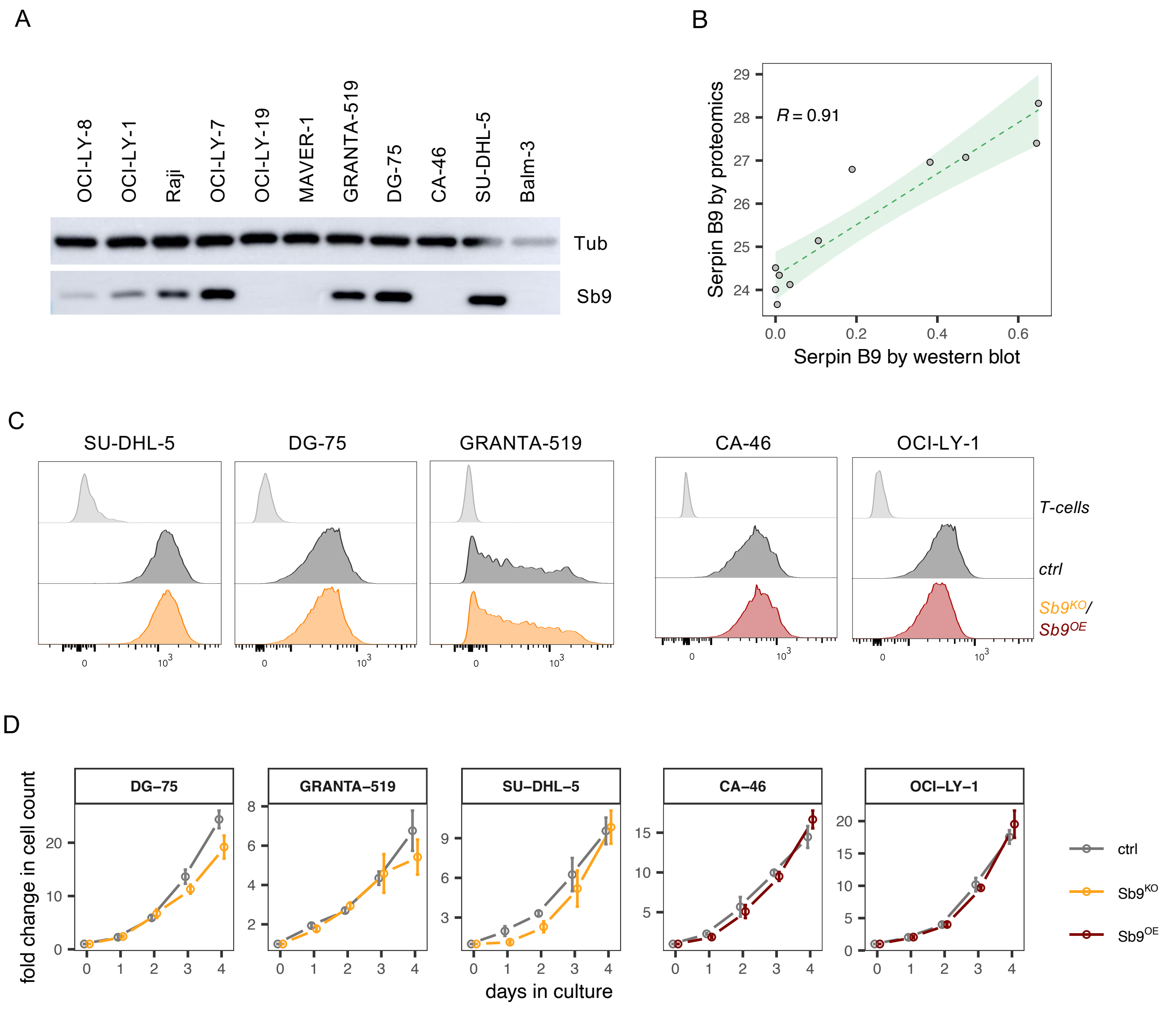


**Supplementary Figure 3. A)** Western blot of of Serpin B9 in selected B-NHL cell lines. **B)** Spearman's correlation of Serpin B9 quantified by proteomics and by western blotting. R denotes the correlation coefficient, the green dashed line denotes the regression line. **C)** CD19 fuorescence intensities measured by flow cytometry in T-cells (negative control) and indicated $Sb9^{KO}$, $Sb9^{KO}$, or control transduced B-NHL cell lines. **D)** Cell growth of indicated $Sb9^{KO}$, $Sb9^{KO}$, or control transduced B-NHL cell lines. Dots indicate means of technical triplicates, bars indicate the standard deviation. Sb9: Serpin B9. Tub: $\alpha$-Tubulin. KO: knock-out. OE: overexpression. ctrl: control.

### Supplementary Figure 4

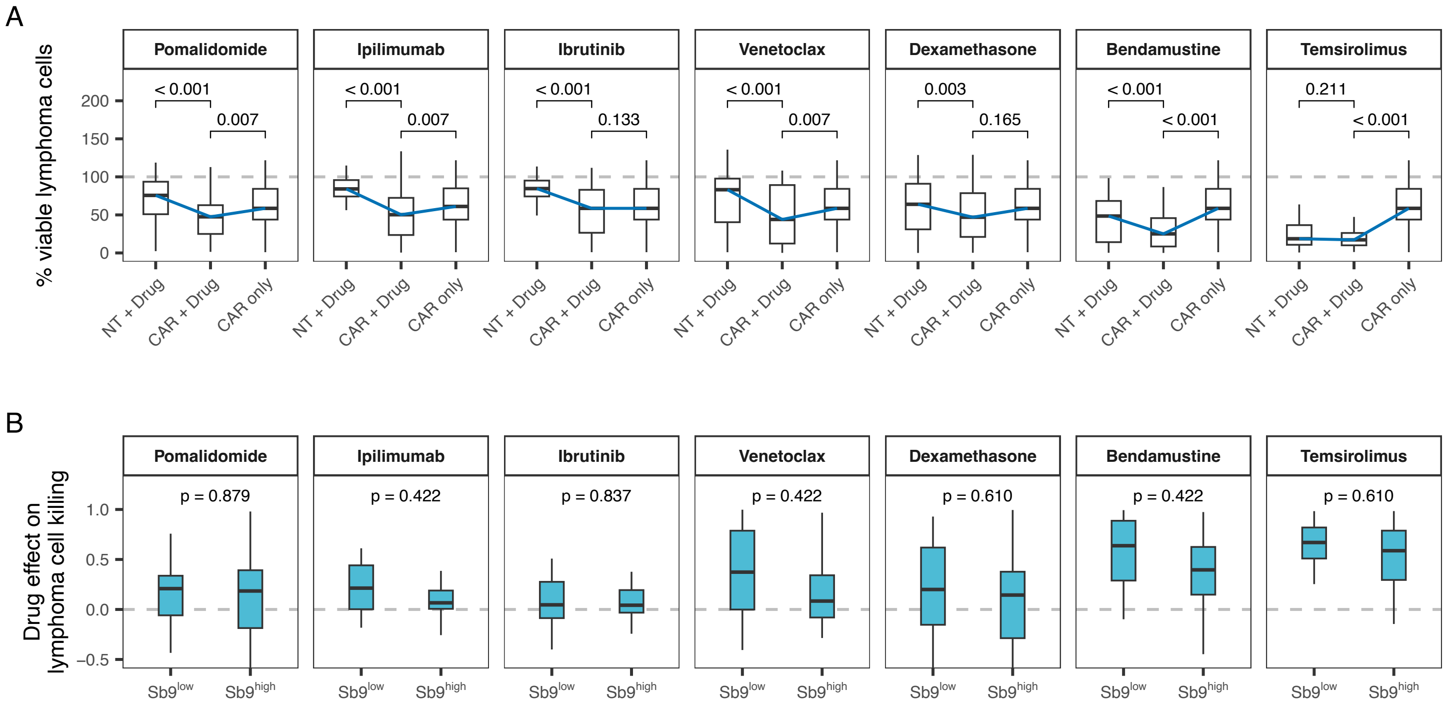


**Supplementary Figure 4. A)** Percentage of viable lymphoma cells after 4-day co-culture with NT or CD19-CAR T-cells with or without a given drug. Blue lines connect the medians of each co-culture/drug-condition to ease interpretation. **B)** Comparison of drug effects on lymphoma cell killing between cell lines with Serpin B9 levels above (Sb9$^{high}$) or below (Sb9$^{low}$) median Serpin B9 abundance. P-values of indicated comparisons were calculated using the two-sided paired (A) or non-paired (B) Wilcoxon's test and are denoted in the plots. Sb9: Serpin B9. NT: non-transduced.

### Supplementary Figure 5

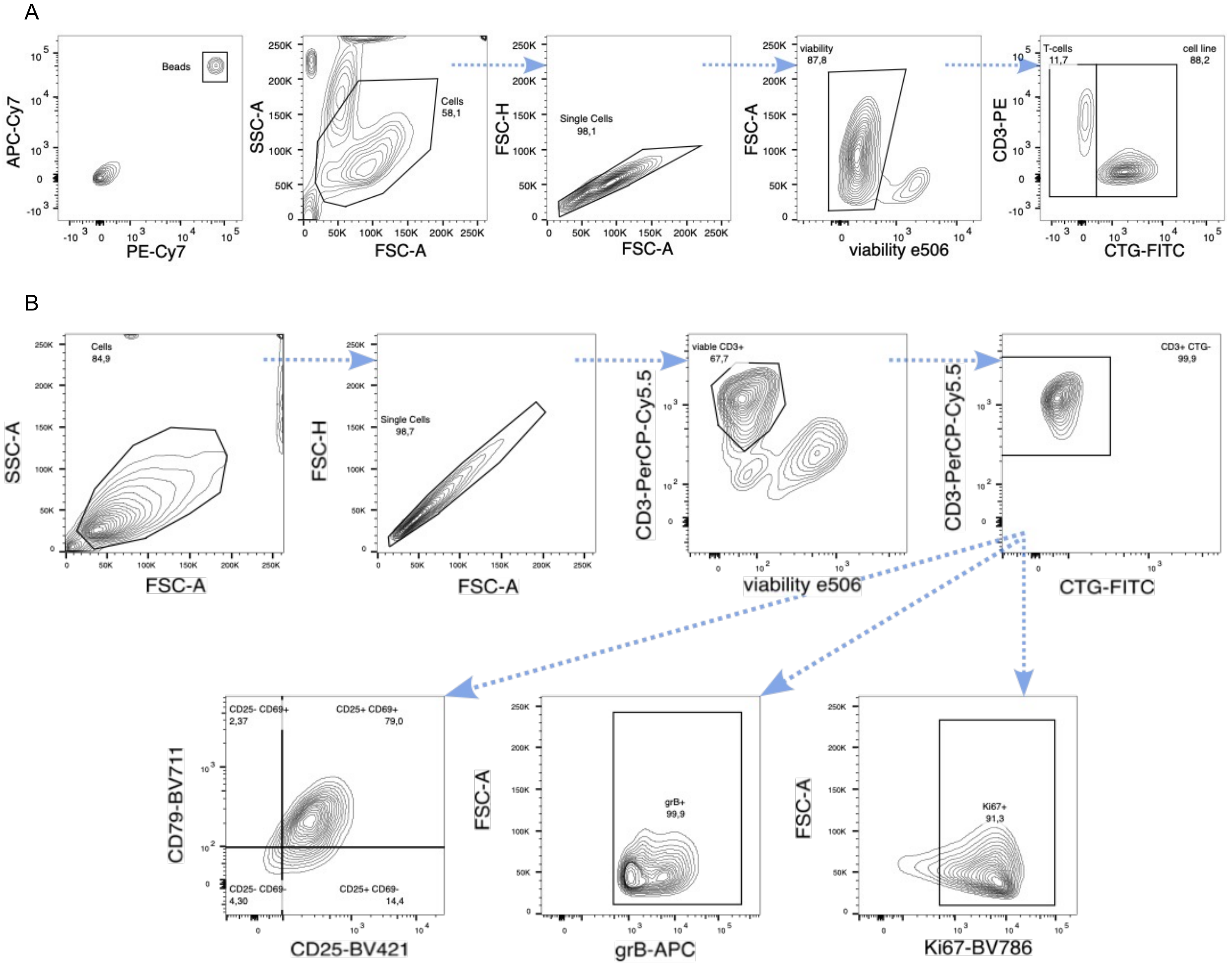


**Supplementary Figure 5. A)** Contour plots illustrating the gating strategy used to identify lymphoma cells and CD19-CAR or NT T-cells within co-cultures after 4-day incubation. **B)** Contour plots illustrating the gating strategy to identify CD25$^{+}$CD69$^{+}$, grB+, and Ki67$^{+}$ viable T-cells in co-cultures with B-NHL cell lines after 3-day incubation. grB: granzyme B. Sb9.
