## Supplementary Methods for "Proteomic profiling identifies Serpin B9 as mediator of resistance to CAR T-cell and bispecific antibody treatment in B-cell lymphoma"

*Proteomics profiling and data processing*

Pellets of $5x10^6$ cells per B-NHL cell line were washed two times with PBS and snap-frozen. Cell pellets lysed by lysis buffer (1 % SDS, 100 mM Hepes/NaOH pH 8.5, and EDTA-free protease inhibitor). Samples were heated to 95 °C for 5 min. DNA and RNA were degraded by the addition of benzonase at 4 °C following incubation for 1 h at 37°C. 10 µg of each lysate were subjected to an in-solution tryptic digest using a modified version of the Single-Pot Solid-Phase-enhanced Sample Preparation (SP3) protocol [1,2]. To this end, lysates were added to Sera-Mag Beads (Thermo Scientific) in 10 µL 15 % formic acid and 30 µL of ethanol. Binding of proteins was achieved by shaking for 15 min at room temperature. SDS was removed by 4 subsequent washes with 200 µL of 70 % ethanol. Proteins were digested overnight at room temperature with 0.4 µg of sequencing grade modified trypsin (Promega) in 40 µL Hepes/NaOH, pH 8.4 in the presence of 1.25 mM TCEP and 5 mM chloroacetamide (Sigma-Aldrich). Beads were separated, washed with 10 µL of an aqueous solution of 2 % DMSO and the combined eluates were dried down.

Peptides were reconstituted in 10 µL of $H_2O$ and reacted for 1 h at room temperature with 80 µg of TMT10plex (Thermo Scientific) [3] label reagent dissolved in 4 µL of acetonitrile. Excess TMT reagent was quenched by the addition of 4 µL of an aqueous 5 % hydroxylamine solution (Sigma). Peptides were reconstituted in 0.1 % formic acid, mixed to achieve a 1:1 ratio across all TMT-channels and purified by a reverse phase clean-up step (OASIS HLB 96-well µElution Plate, Waters).

Peptides were subjected to an offline fractionation under high pH conditions [2]. The resulting 12 fractions were then analyzed by LC-MS/MS on an Orbitrap Fusion Lumos mass spectrometer (Thermo Scientific) as previously described [4]. To this end, peptides were separated using an Ultimate 3000 nano RSLC system (Dionex) equipped with a trapping cartridge (Precolumn C18 PepMap100, 5 mm, 300 µm i.d., 5 µm, 100 Å) and an analytical column (Acclaim PepMap 100. 75 × 50 cm C18, 3 mm, 100 Å) connected to a nanospray-Flex ion source. The peptides were loaded onto the trap column at 30 µL per min using solvent A (0.1 % formic acid) and eluted using a gradient from 2 to 40 % Solvent B (0.1 % formic acid in acetonitrile) over 2 h at 0.3 µL per min (all solvents were of LC-MS grade). The Orbitrap Fusion Lumos was operated in positive ion mode with a spray voltage of 2.4 kV and capillary temperature of 275 °C. Full scan MS spectra with a mass range of 375–1500 m/z were acquired in profile mode using a resolution of 120,000 (maximum fill time of 50 ms or a maximum of $4x10^5$ ions (AGC) and a RF lens setting of 30 %. Fragmentation was triggered for 3 s cycle time for peptide like features with charge states of 2–7 on the MS scan (data-dependent acquisition). Precursors were isolated using the quadrupole with a window of 0.7 m/z and fragmented with a normalized collision energy of 38. Fragment mass spectra were acquired in profile mode and a resolution of 30,000 in profile mode. Maximum fill time was set to 64 ms or an AGC target of $1x10^5$ ions). The dynamic exclusion was set to 45 s.

Acquired data were analyzed using IsobarQuant [5] and Mascot V2.4 (Matrix Science) using a reverse UniProt FASTA Homo sapiens database (UP000005640 from May 2016) including common contaminants. The following modifications were taken into account: Carbamidomethyl (C, fixed), TMT10plex (K, fixed), Acetyl (N-term, variable), Oxidation (M, variable) and TMT10plex (N-term, variable). The mass error tolerance for full scan MS spectra was set to 10 ppm and for MS/MS spectra to 0.02 Da. A maximum of 2 missed cleavages were allowed. A minimum of 2 unique peptides with a peptide length of at least seven amino acids and a false discovery rate below 0.01 were required on the peptide and protein level [6].

The raw output files of IsobarQuant (protein.txt-files) were processed using the R programming language (version 4.2.2). Preprocessing steps were done using the R package matrixQCvis [7] (version 1.4.0). Only proteins that were quantified with at least two unique peptides and present in at least 6 TMT channels were considered for the analysis. Raw signal-sums (signal_sum columns) were normalized using variance stabilization normalization[8] followed by MinDet imputation, setting each missing protein abundance to their 0.01 quantile.

*Protein differential abundance analysis*

Protein differential abundance between cell lines ($n_{samples} = 70$ and proteins = 4873) with respect to their percentage of viable lymphoma cells after treatment with CAR T-cells and BsAb, was assessed using the R package limma [9] (version 3.52.4). To account for potential batch effects TMT channels were included as covariates. P-values were adjusted using the Benjamini-Hochberg procedure [10]. Differentially abundant proteins were defined as adjusted p-value < 0.05.
